# BaMBo: An Annotated Bone Marrow Biopsy Dataset for Segmentation Task

**DOI:** 10.1101/2024.10.02.616393

**Authors:** Anilpreet Singh, Satyender Dharamdasani, Praveen Sharma, Sukrit Gupta

**Author notes:** Joint first authors (Anilpreet Singh, Satyender Dharamdasani).

## Abstract

Bone marrow examination has become increasingly important for the diagnosis and treatment of hematologic and other illnesses. The present methods for analyzing bone marrow biopsy samples involve subjective and inaccurate assessments by visual estimation by pathologists. Thus, there is a need to develop automated tools to assist in the analysis of bone marrow samples. However, there is a lack of publicly available standardized and high-quality datasets that can aid in the research and development of automated tools that can provide consistent and objective measurements. In this paper, we present a comprehensive **B**one **M**arrow **B**i**o**psy (BaMBo) dataset consisting 185 semantic-segmented bone marrow biopsy images, specifically designed for the automated calculation of bone marrow cellularity. Our dataset comprises high-resolution, generalized images of bone marrow biopsies, each annotated with precise semantic segmentation of different haematological components. These components are divided into 4 classes: Bony trabeculae, adipocytes, cellular region and Background (BG). The annotations were performed with the help of two experienced hematopathologists that were supported by state-of-the-art Deep Learning (DL) models and image processing techniques. We then used our dataset to train a custom U-Net based DL model that performs multi-class semantic segmentation of the images (Dice Score: 0.831 *±* 0.099) and predicts the cellularity of these images with an error of 5.9% *±* 8.8%. This shows the applicability of our data for future research in this domain. Our code is available at https://github.com/AI-in-Medicine-IIT-Ropar/BaMbo-Bone-Marrow-Biopsy.

## 1. Background

Bone **marrow** *is* the soft, spongy tissue found inside the center of most bones such as the sternum, pelvis, femur and tibia It’s made up of islands of blood-forming cells called hematopoetic tissue, and fat cells called adipocytes within a network of trabecular bone. It serves as an important part of the hematopoietic system, responsible for the production and maturation of blood cells-erythrocytes, granulocytes, monocytes, lymphocytes and platelets (Travlos, 2006). A bone marrow biopsy is a critical medical procedure where a small amount of bone marrow tissue is removed from the bone for examination under a microscope. Besides aiding in assessment of bone marrow’s ability to produce blood cells, it provides essential diagnostic and prognostic information for a wide range of conditions affecting the bone marrow and blood cells. It helps diagnose various hematological conditions, such as leukemia, lymphoma, multiple myeloma, aplastic anemia, myelodysplastic neoplasm and certain infections (Percival et al., 2017; Keung et al., 2001). It provides valuable information about the stage and progression of diseases affecting the bone marrow. It is essential for minimal residual disease assessment, thereby aiding in monitoring the efficacy of treatments like chemotherapy or radiation therapy. It can also help in uncovering the underlying cause for unexplained symptoms like persistent fever, unexplained anemia, or incidentally detected abnormal blood parameters.

Under the microscope, a pathologist can examine the bone marrow biopsy to assess its health and identify any abnormalities. This involves assessment of various aspects of the tissue, such as the marrow architecture (e.g., cell lineages, vascular or stromal alterations, inflammation, necrosis), cellular configuration, estimation of iron stores, identification of other features such as pigment, infectious agents, proliferative or neoplastic disorders (Brown and Gatter, 1993). The first and crucial step in the morphological examination of bone marrow biopsy is the assessment of the cellularity in order to determine the proportion of area of the marrow space constituted by hematopoietic cells (Bancroft and Gamble, 2008). The whole process is a visual estimation made by the pathologist, that needs training and expertise. It is subjective, error-prone, has high interobserver variability and poor reproducibility. The more precise, point cloud estimation method, is quite labor-intensive and time-consuming, and is therefore, not widely favored for routine use in most diagnostic laboratories.

Computational Pathology, armed with machine learning and artificial intelligence algorithms, holds significant promise for addressing longstanding challenges in this area. Computer vision has helped automate many tasks in histopatheotliogy over the last decade (Cooper et al., 2023). There are existing computer vision models that help automate the identification of various components of a biopsy by segmenting them for easier understanding (Sarkis et al., 2023; Shiffman et al., 2023; Nielsen et al., 2019). This can aid in identifying various haematopoietic elements and thereby, help identify and classify disorders, which would in turn aid the pathologists in easier and faster diagnostics. Central to the development of computational pathology tools is the availability of high-quality datasets that encompass the diverse cellular compositions and pathological conditions observed in bone marrow specimens. For example, Figure 1 shows a WSI and its respective annotation done by a DL model trained on our data, showing how computer vision can be used to segment various parts of a WSI.

**Figure 1.**
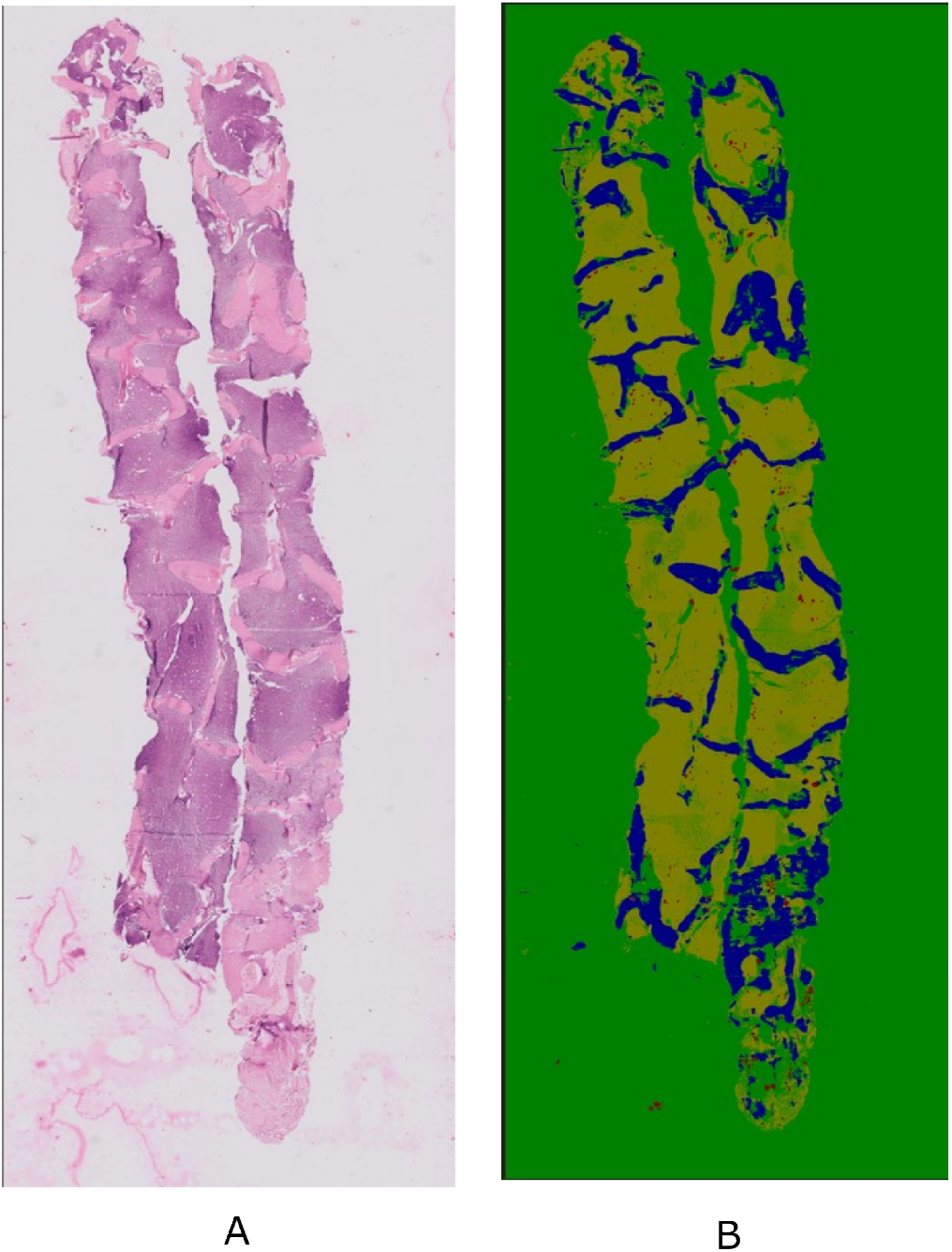
Sample of Whole Slide Image (WSI) of a bone marrow sample collected from a patient at PGIMER. A: Original unlabeled image; and B: Semantic Segmentation performed on the image by our trained DL model. (Colors and the respective classes are: yellow: cells, green: background, blue: bone, red: fat). Credit for the WSI: Dr. Pulkit Rastogi, Department of Histopathology, PGIMER, Chandigarh.

After extensive search our team was unable to find any publicly accessible annotated dataset for bone marrow biopsy images, which hinders the development of opensource diagnostic systems. However, several private annotated datasets have been proposed and used in the literature (van Eekelen et al., 2022; Nielsen et al., 2019; Shiffman et al., 2023; Sarkis et al., 2023). Additionally, none of these studies capture data from Asia or the Indian subcontinent, which has been an under-represented group in digital biomedical data. van Eekelen et al. (2022) annotated PAS-stained bone marrow biopsy images by adding circles on each cell with a diameter equal to the approximated dataset-wide average diameter for that particular cell type. This leads to poor annotations. Shiffman et al. (2023) used private annotated dataset of murine bone marrow biopsy. Paul Cohen et al. (2017) contains a dataset of bone marrow biopsy images, with the center of each cell annotated with a dot, these annotations focus on cell counting. This dataset can prove to be useful for future expansion of our dataset.

It is to be noted that there are open source bone marrow cell identification datasets, that have very high magnification images of individual hematopoeitic cells, such as the ones available on Kaggle and The Cancer Imaging Archive (Matek et al., 2021b; Consortium et al., 2019; Matek et al., 2021a). However, these datasets are different, as they capture the cell morphology on aspirate smears, whereas we present a dataset focusing on Haematoxylin and Eosin (H&E) stained bone marrow biopsies. (Sarkis et al., 2023) have publicly shared 32 WSI of bone marrow biopsies. These, although un-annotated, are very rich in detail and could also be annotated and used as an extension to our dataset.

In this paper, we address the critical gap of non-availability of high quality annotated bone marrow biopsy dataset by presenting a comprehensive dataset of semantic segmented bone marrow biopsy images. We detect the hematopoic region by taking into account blood vessels, tissue cavities caused by specimen shrinkage or tissue fragmentation, large clusters of erythrocytes due to hemorrhage, stain and sectioning artifact and slide preparation faults.

This dataset has four classes viz. Cell, Bone, BG and Fat. These annotations have been meticulously annotated by expert hematopathologists by hand on an annotation platform with the additional help of Machine Learning (ML) models and image processing tools.

## 2. Summary

The BaMBo dataset contains 185 bone marrow biopsy images annotated by expert hematopathologists. It aims to provide an annotated point for researchers building advanced models on bone marrow biopsy images. Since, we provide detailed pixel-wise annotations for the images, the dataset can be used for tasks related to training models for anomaly detection, classification and segmentation. In addition to the pixel-wise classes, we separately provide the area percentage of each class, cellularity, haemorrhage, soft tissue and sectioning artifact. This dataset will not merely serve researchers and clinicians working on the Indian population, but will be a first of its kind public resource globally. We hope that other researchers will use and contribute to this resource. Researchers using their algorithm on our dataset will able to establish the ground truth, even if they do not have an expert hematopathologist onboard, by using the detailed pixel wise class annotations and class metrics made available. The flagging of images containing hemorrhage in our dataset, can aid in developing models for automated detection of hemorrhage in the bone marrow biopsies.

As an application for the BaMBo dataset, we have shown a use case of this dataset for train a DL model for performing segmentation of the bone marrow biopsy image and then used the computed segmentation to compute the cellularity of the images. Evaluating cellularity helps healthcare providers determine the underlying causes of abnormal blood cell counts and formulate appropriate treatment plans. We have provided both the code for our cellularity prediction model https://github.com/AI-in-Medicine-IIT-Ropar/BaMbo-Bone-Marrow-Biop and the dataset in a AI ready format so that it can be readily used by researchers for developing models to solve problems in this domain.

In the subsequent sections of this paper, we delve into resource availability (section 4), the methodology employed for dataset collection and delve into the extensive annotation procedure and methodology and tools employed for it (section 5) and show a use case for training semantic segmentation models and predicting cellularity of bone marrow biopsy images using the BaMbo dataset (section 7).

## 3. Discussion

The presented dataset, composed of semantic-segmented bone marrow biopsy images, offers substantial contributions to the field of hematopathology and the development of automated diagnostic tools. This dataset addresses critical gaps in the availability of high-quality, annotated images essential for the training of machine learning models focused on automated bone marrow diagnostics. While there have been datasets published before on bone marrow (detailed discussion in the supplementary section C), but most of them are private datasets. Despite the strengths of this dataset, there are some limitations to consider. The dataset, while comprehensive lacks segmentation of cellular morphology as different individual cells for eg. erythroid precursors, granulocytes, histiocytes and megakarocytes etc. To develop models that can diagnose diseases by understaning the haematological structure. Blood vessels, plasma and soft tissue can also be segmented into their own unique classes. Future efforts should focus on expanding the dataset to include a broader range of classes and also, to increase the number of annotated images. This additional data can then help in diagnosing bone marrow based diseases at a cellular level.

## 4 Resource Availability

### 4.1 Potential Use Cases

#### Automated Diagnostic Tools and Clinical Decision Support

The dataset can be used to train machine learning models, particularly DL algorithms, to automatically estimate bone marrow cellularity (Section 7). This facilitates faster, more accurate, and standardized assessments compared to the traditional manual microscopic examination, potentially reducing human error and inter-observer variability. This is more precise than visual estimation and faster than point cloud estimation. Automated tools developed using the dataset can serve as decision support systems for hematologists and pathologists, helping them diagnose and monitor various hematological condistiyons such as anemia, leukemia, and myeloproliferative disorders. These tools can provide non-subjective and quantitative data that support clinical decisions regarding treatment strategies and disease monitoring.

#### Research in Hematopathology

From a research perspective, this dataset enables future in-depth studies of the relationships between bone marrow cellularity and various hematological diseases. By providing detailed annotations, the dataset supports the identification and precise structural analysis of pathological features within bone marrow biopsies. This can lead to a better understanding of disease mechanisms and the development of new therapeutic approaches.

#### Educational Applications

In the realm of education, this dataset serves as a valuable training resource for medical students and residents. The segmentation provides clear visual insights into the cellular architecture of bone marrow, enhancing the learning experience with real-world examples. This can improve the understanding of the structural and visual differences between features in bone marrow pathology and the nuances of artefactual changes such as procedural hemorrhage, sectioning tears, tissue retraction, etc.

### 4.2 Licensing

The dataset will be hosted on an open-source platform for access by public. It may be obtained by submission of an online application form and acceptance of a Data Use Agreement. The application must include the investigator’s institutional affiliation and the proposed uses of the BaMBo dataset.

### 4.3 Ethical Considerations

All procedures performed in studies involving human participants were per the ethical standards of the institutional and/or national research committee and with the 1964 Helsinki Declaration and its later amendments or comparable ethical standards. The Institutional Ethical Clearance Committee (IEC) approval was obtained for the conduct of the study.

Subjects’ consent was taken for using their biopsy images for research purposes. Subjects were verbally (in their vernacular language) informed about the option to opt out of the study and only the data for those subjects who opted in was included in the study. No extra samples were obtained from the patients. Data was obtained from the images of Hematoxylin and Eosin (H & E) stained bone marrow biopsy slides, which is a part of routine diagnostics of various disease conditions.

## 5 Methods

Our team used machine learning to help pathologists with the annotation process. We processed the images using Segment Anything Model (SAM) segmentation model which produced segmentations of the parts of the biopsy image. The visual characteristics and patterns of objects in each class was understood carefully to extract these features into a custom feature vector, to classify every object that SAM segmented, k-means and Support Vector Machine (SVM) models were trained through active learning, by initially training using manually classified data and then using corrected outputs. This pipeline of automatic annotation has made our work very efficient and has also been since used for development of other datasets in our lab. These annotations were of good quality but needed expert refinement, Pathologists at PGIMER, Chandigarh used remote access to extensively refine annotations of these images. Careful annotations of 185 images took 15-20 minutes for each image and over 2 months. These annotations are in the integer-encoded data format which can be easily used to train machine learning models. Along with the annotations, a data table was also made that includes the details of each image i.e. percentage of cell, bone, fat, BG, cellularity and whether each image has hemorrhage, soft tissue or sectioning artifact.

### 5.1 Data Collection

Data collection involved the acquisition of Bone Marrow Trephine (BMT) biopsy specimens using standardized procedures at Postgraduate Institute of Medical Education and Research, Chandigarh (PGIMER). These procedures, as described later, are medical standards as mentioned in (Bancroft and Gamble, 2008). The dataset comprises 185 biopsy images from a total of 38 patients, diagnosed with varied hematological states covering myeloproliferative neoplasms, leukemia, aplastic anemia, uninvolved staging marrows among others. Subjects were chosen at random from both the genders, contains a large age gap and varied medical history. All the patient related identifiers were removed from the dataset and the dataset was completely anonymized for patient confidentiality. Ethics were kept in consideration as described in Section 4.3.

Biopsis are taken from the posterior superior iliac spine. During the procedure, a larger needle was employed to remove a small core of the marrow tissue (biopsy) (Cloos et al., 2018; Rindy and Chambers, 2020). Biopsy samples then underwent Decalcification, Fixation and Paraffin embedding. They were then prepared for microscopic examination by sectioning into 2-3 microns samples and staining with Hematoxylin and Eosin (H&E). High-resolution digital images of stained biopsy sections were captured using an Olympus BX53 microscope at a magnification of 100x, 200x and 400x.

The data collection was conducted over a period of a three months and resulted in around 300 images. Out of the 300 bone marrow biopsy images obtained from PGIMER, images with poor clarity and repetitive images were excluded. After thorough inspection, 185 high quality images were retained. Thorough analysis was done on these images. The characteristics of the BaMbo dataset are summarized in Table 1. The Cellularity range of all the images, (Cellularity explained in Section 7), was plotted in a bar graph with a 10% interval on the x-axis, effectively illustrating the distribution in (Figure 2).

**Table 1:** Characteristics of BaMbo Dataset.

| Quantity | Value |
| --- | --- |
| Total Number of Images | 185 |
| No. of Patients | 38 |
| Age (Range) | 4-65 Years |
| Gender | 20 Males |
| Image Resolution | 1200 × 1920 |
| Images with haemorrhage | 40 |
| Images with Artifact/Tears | 18 |
| Images with Soft Tissue | 5 |
| Cellularity | 56.23 ± 37.39% |

**Figure 2.**
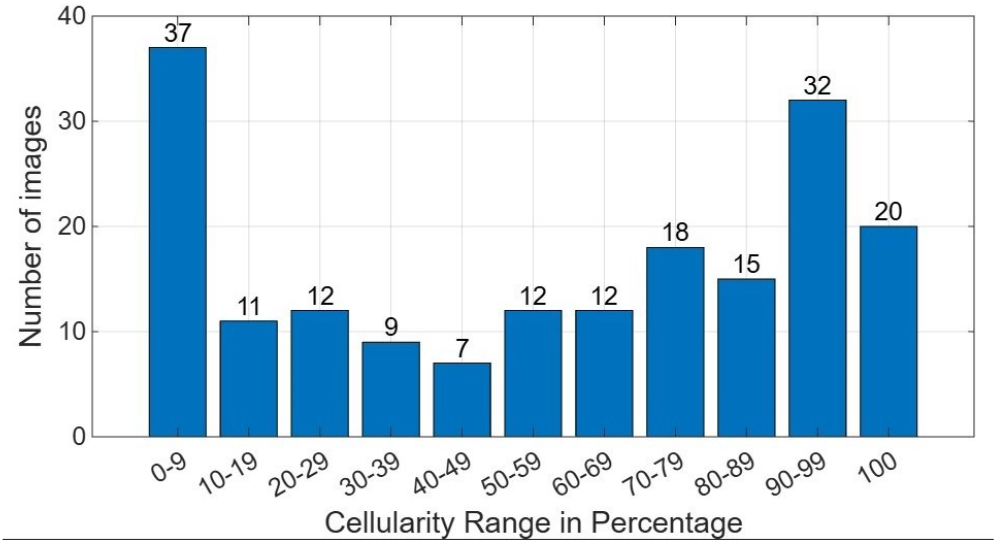
Distribution of Cellularity among images in BaMBo Dataset

## 6 Data annotations

In order to perform the annotation of our biopsy specimens, we developed a preprocessing pipeline for first generating a rough annotation of the images. This pipeline made the annotation process faster and more efficient as the pathologist did not need to start from scratch. For this pipeline, we utilized a state-of-the-art segmentation model SAM (Kirillov et al., 2023) and the Computer Vision Annotation Tool (CVAT) platform for final annotation by the pathologist. SAM is a promptable segmentation system with zero-shot generalization to unfamiliar objects and images, without the need for additional training. (Figure 3) illustrates the process of generating automatic annotations. More information on the annotation format is in the Supplementary (Section A).

**Figure 3.**
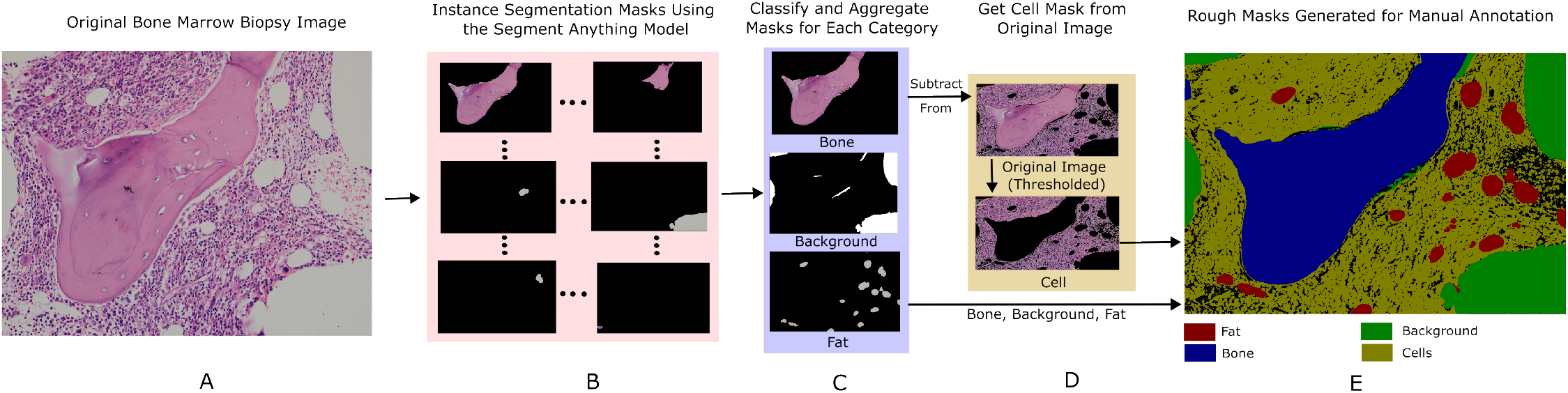
Flowchart representation of generating rough automated annotations by SAM, SVM and image processing. A: Original BMT image. B: Generating unclassified segmentation objects using SAM on basis of visual distinction betwen various parts of image. C: Classification of segmentation objects into masks by SVM on the basis of feature vector of colors,texture and shape. D: Generation of cell mask by first using color thresholding to seperate non-white part from original image and subtracting bone mask from it to obtain aproximate cell mask. E: Stitching together all 4 masks and obtaining rough annotations.

### 6.1 Obtaining Automated Rough Annotations

#### 6.1.1 SAM Segmentation

The SAM performs instance segmentation in images, and is known to be robust to the presence of unfamiliar objects or complex backgrounds. This enables SAM to identify and delineate distinct objects or regions within the image. We used SAM to first perform instance segmentation on the entire image thereby transforming it into distinct, non-overlapping segmentation objects. These segmentation objects contain parts of the biopsy detached from the complete image based on its distinctive appearance to its surroundings. Hyperparameters used for initilizing SAM have been detailed in the Supplementary (Section B).

#### 6.1.2 Classification of Segmentation Objects Generated by SAM

Processing SAM on bone biopsy images gave distinct, non-overlapping and uncategorized segmentation objects for bone, fat and backgorund (bg). However since the cellular regions in the image were very dense, SAM segmented individual cells instead of taking the complete cellular region into one area, which generated multiple small segmentation objects for each cell. For each image, all the generated segmentation objects were unclassified and needed to be classified into one of the 4 classes viz. cell, bone, fat and bg.

Since, there were no pre-existing datasets or models to perform the classification of segmentation objects, we manually classified a subset of these objects by observing visual differences between segmentation objects of each class. Fats are white eliptical structures, bones are pinkish and large in size, cells are small and very dense closely packed structures with purple (nucleus) and pink color (cytoplasm), and the slide bg is white and of non-uniform shape. Using image processing techniques, we extracted feature vector from each segmentation object (Feature vector detailed in Supplementary Section B). For the first subset (3681 segmentation objects of 51 images), we first used K-means clustering to cluster the feature vector of these objects into 4 classes. This was further manually checked and corrected. This formed our ground truth database for further segmentation object classification and was used to train a SVM model to perform classification of unknown segmentation objects. We performed classification for another batch of 40 images and manually checked the assigned classes to augment our training set further. We iterated this procedure till we generated the test results for the final set of images. The final SVM model had 96% segmentation object classification accuracy.

#### 6.1.3 Obtaining the segmentation objects for cellular regions

In the last step, we have categorized segmentation objects of bone, fat and the bg. However, as described earlier, the segmentation objects for the cellular regions were of low quality and therefore had to be discarded. We noted that only fat and bg have completely white segmentation objects and therefore the only non-white regions left belong to either cells or bones. We, therefore, used color thresholding set to obtain the non-white region of the image and further removed the bone segmentation objects from the obtained region, thereby giving us the combined segmentation object for cells. The obtained cell objects also do not have any unwanted white region that might have been small parts of fat or bg. It is to be noted that in this process some parts that should have been cells such as blood vessels, were not being included due to their white color, therefore cell masks had much refinement to be done.

### 6.2 Manual annotation by Pathologists

The automated annotation pipeline gave us a rough annotation of each image. It is important to note that these annotations were not devoid of inaccuracies and that this was expected. These inaccuracies included poor segmentation of cell boundaries, blood arterioles were misclassified as BG or fat instead of cells, some irregular adipocytes were misclassified as BG, soft tissue and artifacts were misclassified. The goal of the above exercise was to give a headstart to the pathologists, thus decreasing the time taken for the annotation process. The images with their automated segmentation objects were given to expert hematopathologists to resolve the issues with the segmentation.

(Figure 4) illustrates the manual annotation procedure, In the first step of the procedure the automated annotations were converted into PascalVOC format and uploaded to CVAT (Sekachev et al., 2020). It is an open-source software used for image and video annotation tasks, it was chosen as our annotation platform due to its intuitive userinterface and open-source nature. Each image was then thoroughly annotated by an expert. Our main concern in annotating biopsy was accurate annotation of various components into respective categories of cells, fat, bone and BG. The images were annotated over the course of multiple sessions for different batch of images. Each image took around 15-20 minutes to annotate by a hematopathologist and it took 2 months to complete all annotations. The characteristics of the images after annotation in given in the Table 2.

**Table 2:**
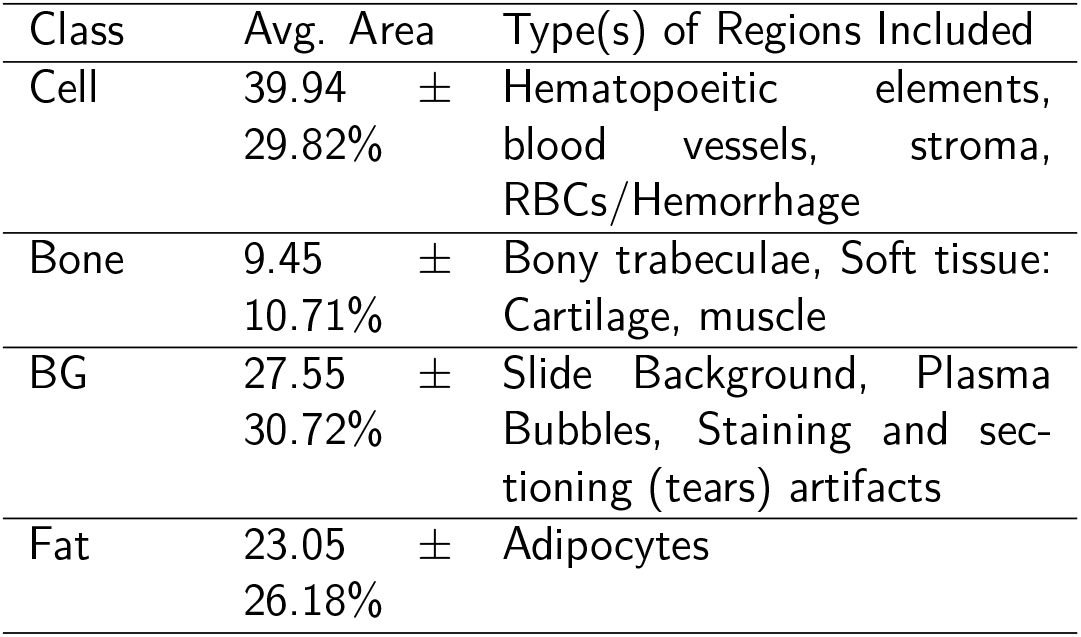
Description of the different classes in the training dataset with their areas and types of regions covered by them.

| Class | Avg. Area | Type(s) of Regions Included |
| --- | --- | --- |
| Cell | 39.94 ± 29.82% | Hematopoietic elements, blood vessels, stroma, RBCs/Hemorrhage |
| Bone | 9.45 ± 10.71% | Bony trabeculae, Soft tissue: Cartilage, muscle |
| BG | 27.55 ± 30.72% | Slide Background, Plasma Bubbles, Staining and sectioning (tears) artifacts |
| Fat | 23.05 ± 26.18% | Adipocytes |

**Figure 4.**
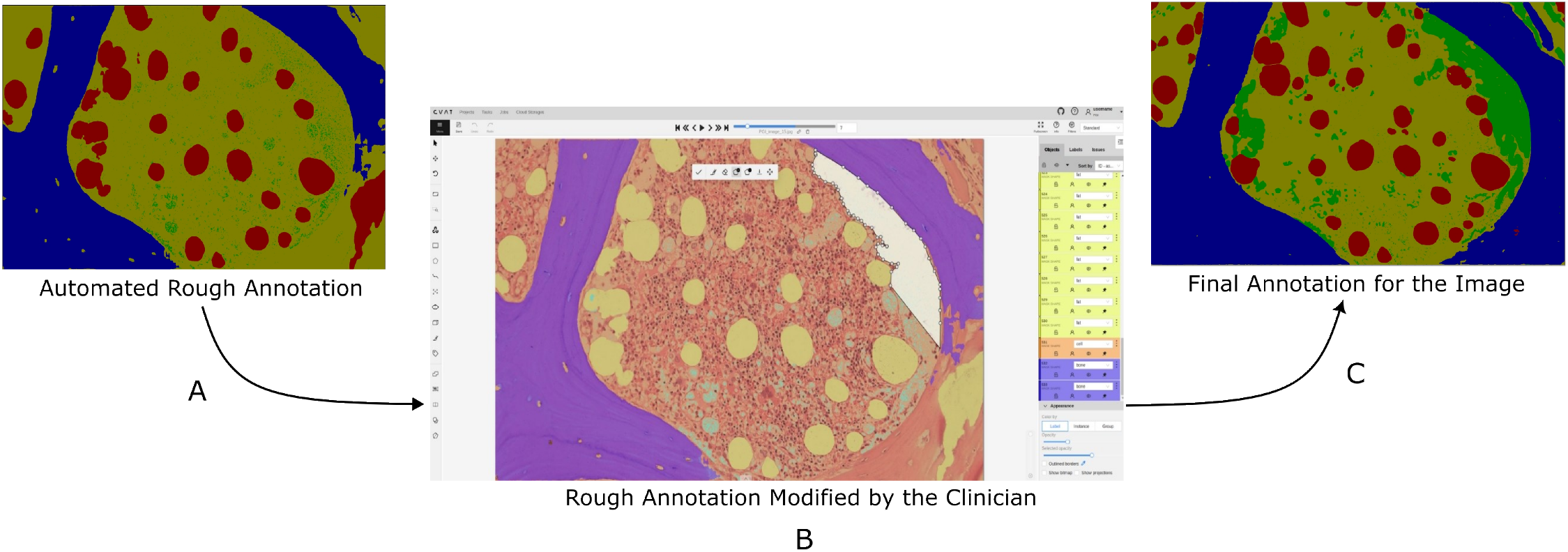
Procedure of annotation by experts. A): Automated rough annotations from automated annotation pipeline taken as input. B): Pathologist using CVAT remotely to precisely edit the annotation to refine it. C): Final precise annotation to be taken as ground truth (Colors and the respective classes are: yellow: cells, green: background, blue: bone, red: fat)

## 7 Using the BaMBo Dataset for Cellularity Prediction

### 7.1 Motivation

Cellularity is an important characteristic of any bone marrow biopsy diagnosis, it describes the proportion of hematopoietic (blood-forming) cells present in the bone marrow. This measure is crucial for evaluating the bone marrow’s health and function. Aberrations in bone marrow cellularity frequently serves as a biomarkers of underlying pathologic processes, encompassing both hematologic malignancies and systemic disorders that demonstrably impact hematopoiesis. High cellularity may indicate conditions such as leukemia or myeloproliferative disorders, where there is an abnormal increase in blood-forming cells (Nielsen et al., 2019). Low cellularity could suggest bone marrow failure conditions, such as aplastic anemia, where there is an inadequate production of blood cells. Thus, evaluating cellularity helps healthcare providers determine the underlying causes of abnormal blood cell counts and formulate appropriate treatment plans. Cellularity of any patient can be calculated by observing the biopsy under microscope. Cellularity is calculated as:

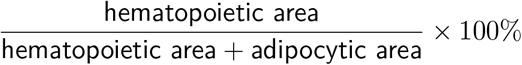

Currently pathologists manually estimate the cellularity percentage of a bone marrow biopsy by visualizing it under the microscope. This method besides being subjective, it is error-prone with poor reliability and high inter-observer variability. We, therefore, aim to train a model that will perform semantic segmentation of the bone marrow biopsy image into different classes and can thus help in computing a reliable, objective, reproducible and more accurate estimate of the cellularity.

### 7.2 Dataset Preparation

While preparing the dataset, we made sure that there is no leakage of images from the train set to the test set. Therefore, we labeled an additional set of 44 images from 9 subjects for the test set. We got 183 images in the training set and 44 images in the test set. After brightness and flip augmentation total images in training set were doubled to 366, no augmentation was applied on testing.

### 7.3 Model Used

We performed these simulations to show that off-the-shelf models that can be used on top of our datasets to perform meaningful tasks that lead to reduction of clinical efforts and give accurate performance. Our focus was on training model architectures suitable for the tasks at hand (semantic segmentation, here) instead of optimizing model performances.

A U-net model with pretrained encoder on ImageNet dataset and a custom decoder had 12 residual blocks and 4 skip connections. The architecture of the decoder was taken from Kaggle. For the model training all the encoder layers were freezed and only the decoder layers were trained. We used a learning rate of 10^*−*4^ with an AdamW optimizer, a batch size of 8, the maximum number of epochs to 140 and early stopping with a patience of 10. Dice loss (1 -Dice Coefficient) was taken as the loss function, while dice coefficient and Mean IoU were also used as metrics for model performance. Our models were trained using 5 random seeds and implemented using the Tensorflow Python library (TensorFlow, 2018)and run on a NVIDIA T1000 8GB GPU. The model trained on 200 epochs and had a uniform decrease in loss and uniform increase in both dice coefficent and mean IoU.

### 7.4 Results

mIOU and Dice Coefficient for the training and testing set w.r.t. epochs is plotted in the Supplementary section E.

#### 7.4.1 Segmentation

The model has a dice coefficient score of 0.8319 *±* 0.099 and a mean IoU score of 0.7122 *±* 0.1443. The smooth training and testing loss curves (Figure 6) and testing evaluations metrics in (Table 3) shows how the Xception model was able to accurately fit to the data. (Figure 5) is the results of the inference on one of the test images. These performance metrics prove that the model can be successfully used for segmentation of various parts of a biopsy for further diagnostics and other models.

**Table 3:** Description of model performance across five different random initializations. Abbreviations: Mean Absolute Error (MAE) and Mean Absolute Percentage Error (MAPE).

| Task | Metric | Mean ± Std. Dev. |
| --- | --- | --- |
| Segmentation | Dice | $0.83 \pm 0.09$ |
| | mIoU | $0.71 \pm 0.14$ |
| Cellularity | MAE | $0.059 \pm 0.088$ |
| | MAPE | $5.9\% \pm 8.8\%$ |

**Figure 5.**
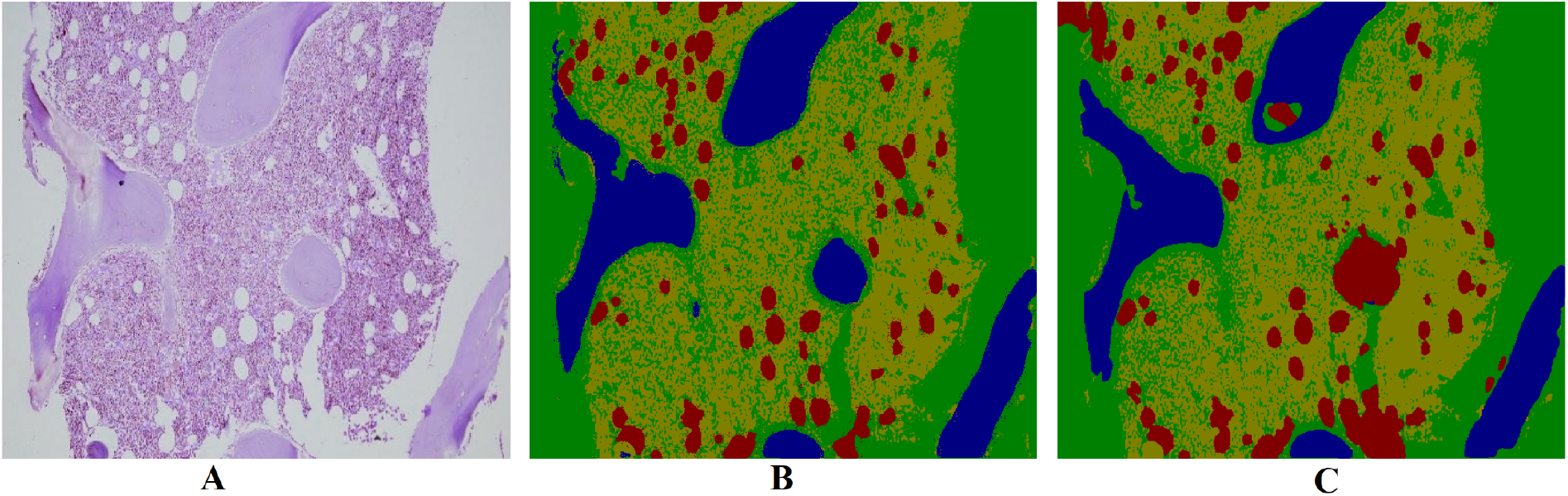
(A) Sample image from the test set (B) Image with ground truth masks overlaid on top; and (C) image with predicted segmentation masks from our model. (yellow: cells, green: background, blue: bone, red: fat)

**Figure 6.**
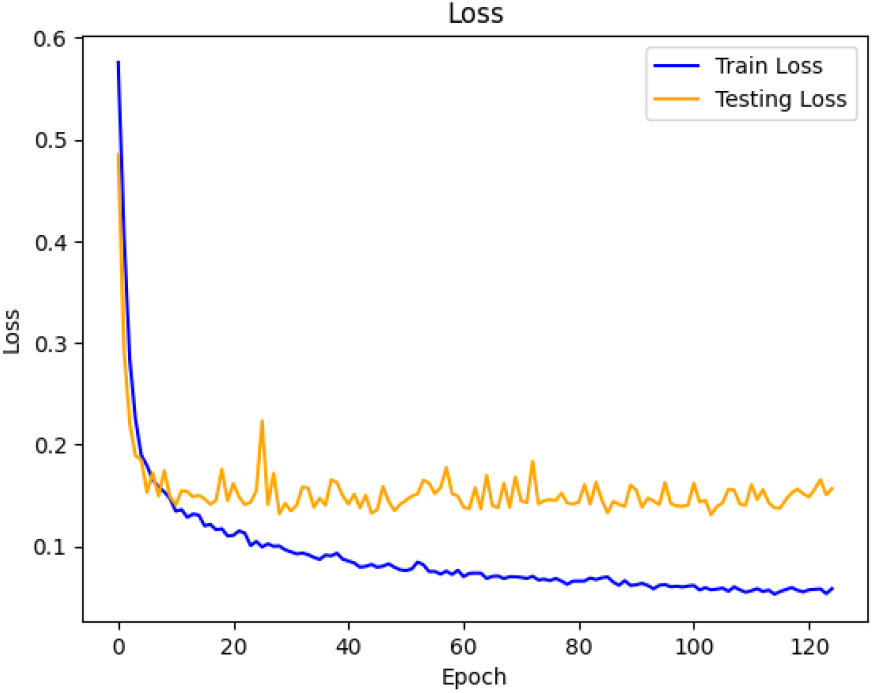
Plot of the dice loss over the number of epochs for the train and test sets for the segmentation model.

#### 7.4.2 Cellularity Percentage Prediction

We computed cellularity by taking into consideration the area of the predicted cell mask and area of the predicted fat mask, using the general formula (described in section 5). The model gives a Mean Absolute Percentage Error of 3.4% while calculating cellularity from fat and cell prediction masks. This means that using the segmentation masks obtained from our segmentation model, we were able to estimate cellularity with a high accuracy establishing the utility of the BaMBo dataset.

## Acknowledgments

We acknowledge important conversations with our colleagues Dr. Pulkit Rastogi, Amit Kumar and Maninder Kaur.

## Ethical Standards

The work follows appropriate ethical standards in conducting research and writing the manuscript, following all applicable laws and regulations regarding treatment of animals or human subjects.

## Data availability

The dataset can be found on an open-source platform for public access. We followed the rules and regulations as mentioned in Starmans and Tsirikoglou (2024) and uploaded the dataset on Singh et al. (2024). To obtain the dataset, users must submit an online application form and accept the Data Use Agreement. The application must include the investigator’s institutional affiliation and the proposed uses of the BoMBR dataset.

