## Supplementary Materials for "BaMBo: An Annotated Bone Marrow Biopsy Dataset for Segmentation Task"

October 3, 2024

#### A Annotation format and cellularity

The dataset is provided in Integer Encoded format. The dataset is organized into subject folders, with each subject folder containing a scans folder. Inside the scans folder, you will find image folders, each of which contains the original image as well as its corresponding annotation. The Integer Encoded masks are one hot encoded images, with pixel having one of 4 pixel values 0,1,2,3, where 0 is bg, 1 is fat, 2 is cell and 4 represents bone. We have also included publish1.csv file that include analysis of each one of the 185 images, such as cellularity and area covered by each class, number of adipocytes and it also includes which images have hemorrhage, soft tissues and sectioning artifacts. You can find the file on the following link: [Publish1.csv]

#### B Hyperparameters

The following are the hyperparamters used for SAM to obtain the segmentation objects:
$sam_checkpoint = "sam\_vit\_h\_4b8939.pth"$
$model_type = "vit\_h"$
$points\_per\_side = 160,$
$pred\_iou\_thresh = 0.85,$
$stability\_score\_thresh = 0.85,$
$min_mask_region_area = 100,$
$stability\_score\_offset = 1.0,$
$box\_nms\_thresh = 0.7$

Each segmentation object was turned to a feature vector for classification by the SVM model, the following was the structure of the feature vector used:

1. Percentage of pink in the segmentation object.
2. Percentage of purple in the segmentation object.
3. Percentage of white in the segmentation object.
4. Circularity of the segmentation Object $\times$ Percentage of white in the segmentation object.
5. Variance in colors in the red channel.
6. Variance in colors in the green channel.
7. Variance in colors in the blue channel.

CVAT (server version 2.11.0) was locally installed on our system (Ubuntu 22.04 x86_64 ), Pathologist from PGIMER, Chandigarh used remote desktop access to our system using AnyDesk and TeamViewer to remotely annotate the dataset. The refined dataset was downloaded in PascalVOC format and then converted to Integer-encoded data with attributes as mentioned above.

### C Related Works

#### C.1 MarrowQuant 2.0: a digital pathology workflow assisting bone marrow evaluation in experimental and clinical hematology

It is a digital hematopathology workflow integrated within QuPath software, which serves as BM quantifier for 5 mutually exclusive compartments (bone, hematopoietic, adipocytic, and interstitial/microvasculature areas and other). For the training of this model 36 BM trephine biopsies were selected from the Institute of Pathology biobank at CHUV. These biopsies were collected for clinical purposes from patients undergoing treatment for AML or MDS at diagnosis and different times after induction chemotherapy. 32 out of these 36 Whole Slide images are available at `https://idr.openmicroscopy.org/webclient/?show=project-2103`. A supplementary csv file is also available with details of each image along with its cellularity and area covered by each class.

#### C.2 Using deep learning for quantification of cellularity and cell lineages in bone marrow biopsies and comparison to normal age-related variation

157 bone marrow trephine biopsies performed for staging of lymphoid or solid cancers in the period 2017–2020 were selected from the archive of the Radboud University Medical Center. Twenty-one randomly selected WSIs were used for the development and evaluation of the neural network, split in a training, tuning, and test set of 14, 5, and 2 WSIs, respectively. The tuning set was used to tune the hyperparameters of the neural network and monitor for overfitting during training. In total, 7864 annotations were made across the training and tuning set. The network weights that performed best on the tuning set were applied as a final model on the test set. The full annotation for the test consisted of nine bounding boxes (on average $500\times500$ $\mu$m) across the two WSIs with a total number of 11,444 annotations.

#### C.3 Automatic bone marrow cellularity estimation in H&E stained whole slide images

Eight WSI H&E stains of bone marrow biopsies from eight different subjects obtained from Aalborg University Hospital, Denmark, were available for the study. An experienced hematopathologist visually inspected the WSI H&E stains to provide an approximate average cellularity estimate within each WSI H&E stain. In each WSI H&E stain, 1–2 subimages ($500 \times 500$ pixels) containing representative bone marrow tissue types were manually extracted for training and validation of the algorithm (13 in total). In each subimage, manual segmentations of the following tissue types were performed: RBM, YBM, erythrocytes, trabecular bone, fibrosis, and fibrin. The manual segmentations were verified by an experienced hematopathologist.

#### C.4 Analysis of cellularity in H&E-stained rat bone marrow tissue via deep learning

For the GT collection, they used male and female rat bone marrow sternum slides from the Genentech study archive, chosen to represent a range of levels of cell depletion. A computational scientist and a technician annotated, on 1 sternebra section per WSI, a boundary around the sternebra, marrow, cartilage, and admixed adipose tissue. Three pathologists and 2 technicians marked all MKC boundaries within the marrow region in 1 sternebra section per WSI. Four pathologists annotated boundaries around small hematopoietic cells in 2 predefined rectangular regions per WSI.

#### D Example Annotations

In the following examples, the first image is the original image while the second image is its corresponding annotation performed by experts. Yellow pixels depict cells, blue pixels depict bone, red pixels depict fat, and green pixels depict background. These images are taken directly from the samples in our published dataset.

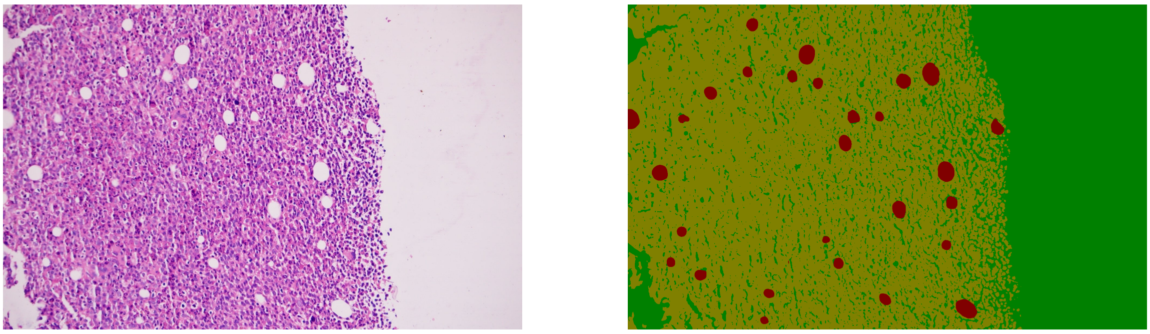

Figure S1: Example 1

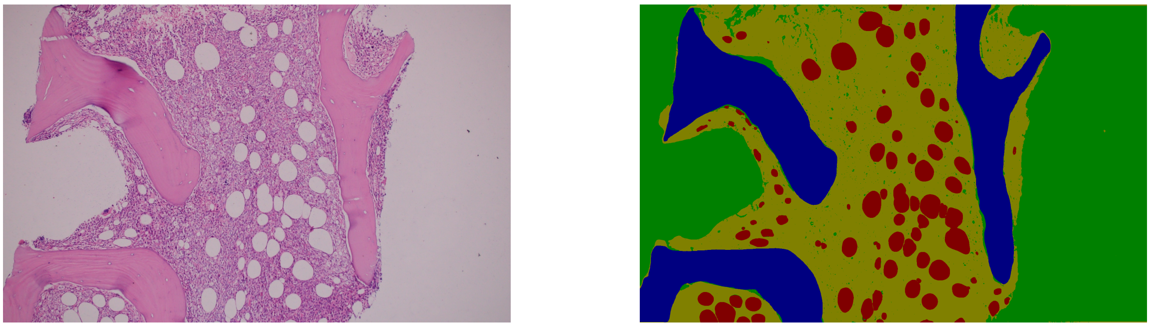

Figure S2: Example 2

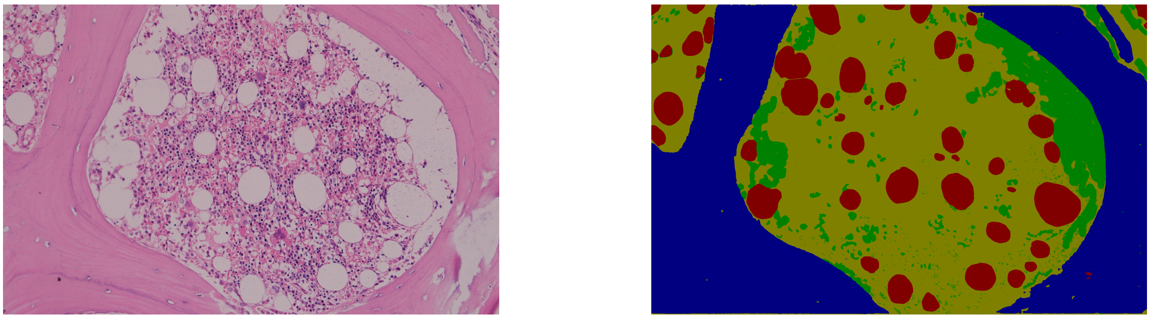

Figure S3: Example 3

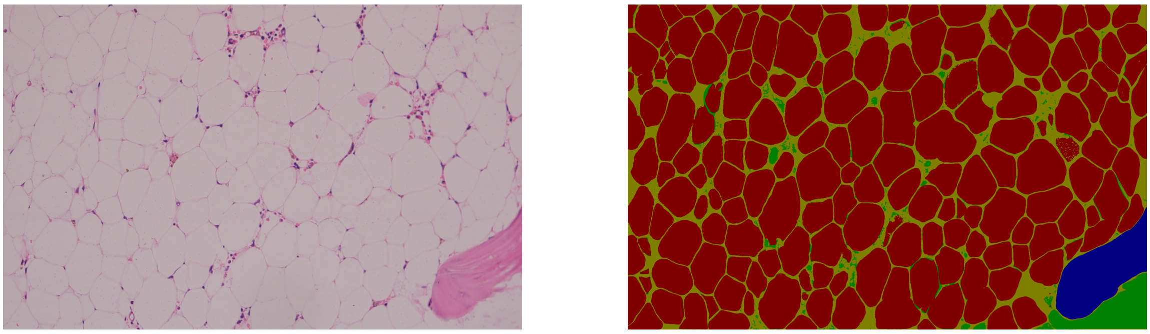

Figure S4: Example 4

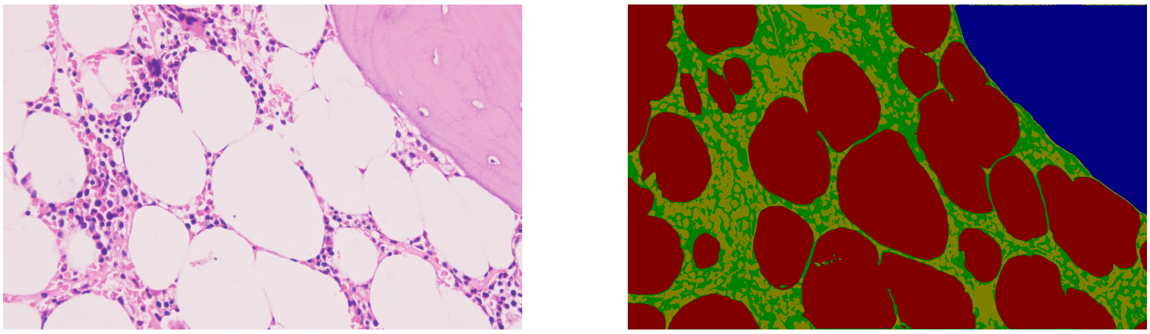

Figure S5: Example 5

#### E Results

The following graphs show the curves for the 109 epochs the model trained on before early stopping. Both training and testing sets were evaluated and plotted, the first graph shows the mIOU score evaluated on Train and Test set at 109 epochs, the second graph shows the Dice coefficient score evaluated on Train and Test set at 109 epochs. It can be observed that the testing performance has converged over the epochs.

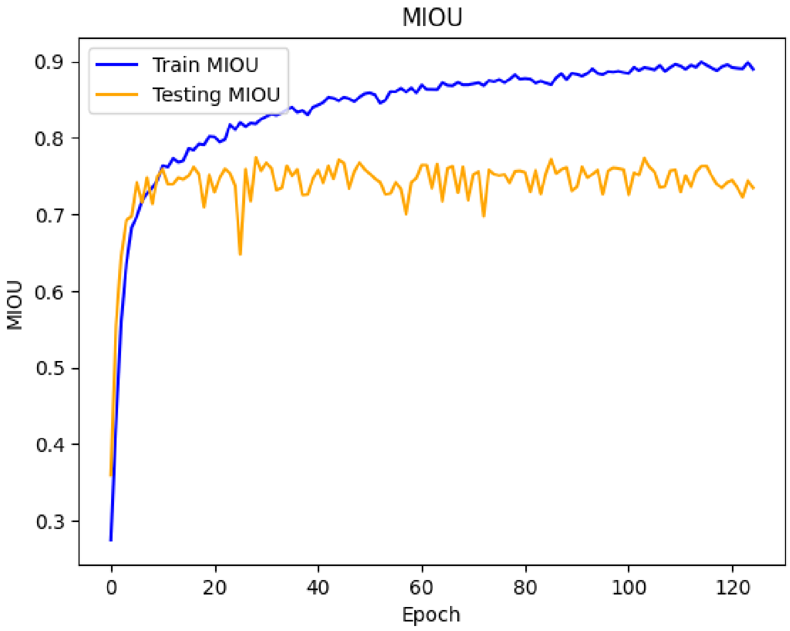


Figure S6: Training and testing mIoU change with training epochs

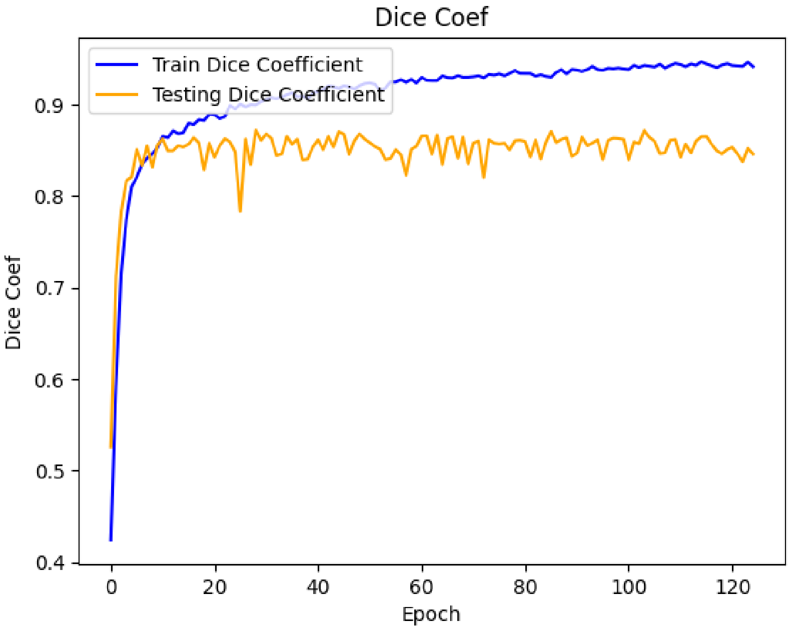


Figure S7: Training and testing Dice Score change with training epochs
